# Transwell-based microphysiological platform for high-resolution imaging of airway tissues

**DOI:** 10.1101/2023.11.22.567838

**Authors:** Amanzhol Kurmashev, Julia A. Boos, Benoît-Joseph Laventie, A. Leoni Swart, Rosmarie Sütterlin, Tina Junne, Urs Jenal, Andreas Hierlemann

## Abstract

Transwell-based airway models have become increasingly important to study the effects of respiratory diseases and drug treatment at the air-liquid interface of the lung epithelial barrier. However, the underlying mechanisms at tissue and cell level often remain unclear, as transwell inserts feature limited live-cell imaging compatibility. Here, we report on a novel microphysiological platform for the cultivation of transwell-based lung tissues providing the possibility to alternate between air-liquid and liquid-liquid interfaces. While the air-liquid interface recapitulates physiological conditions for the lung model, the liquid-liquid interface enables live-imaging of the tissue at high spatiotemporal resolution. The plastics-based microfluidic platform enables insertion and recuperation of the transwell inserts, which allows for tissue cultivation and analysis under standardized well plate conditions. We used the device to monitor infections of *Pseudomonas aeruginosa* in human stem-cell-derived bronchial epithelial tissue. We continuously imaged the progression of a *P. aeruginosa* infection in real time at high resolution, which provided insights into bacterial spreading and invasion on the apical tissue surface, as well as insights into tissue breaching and destruction over time. The airway tissue culture system is a powerful tool to visualize and elucidate key processes of developing respiratory diseases and to facilitate drug testing and development.

## 1. Introduction

*In vitro* models of the human airway have gained increasing importance for studying fundamental aspects of respiratory biology and diseases [1, 2]. Due to their low cost, easy accessibility, and high physiological relevance, *in vitro* human airway models offer promising advantages over commonly used *in vivo* testing with animal models [3]. Particularly, advancements in stem cell technology now allow for a faithful recapitulation of key morphological and functional aspects of the lung airway system *in vitro*, including bronchial and alveolar structures [2]. Bronchial airway tissues, derived from pluripotent stem cells or lung-specific adult stem cells, yielded polarized, pseudostratified epithelial tissue exhibiting *in vivo*-like characteristics, such as a viscous mucus layer covering the epithelium and active cilia beating, mimicking mucociliary clearance.

Along with this development, the use of transwell inserts has become increasingly popular to cultivate epithelial barrier structures *in vitro*. In these systems, airway tissues are cultured on a semipermeable membrane that enables the creation of a physiologically relevant air-liquid interface, establishing transwells as standard tools for permeability and transport studies across the epithelial barrier [4–8]. Moreover, transwells are easy to handle and well-suited for high-throughput applications due to their availability in various multi-well plate formats. As such, transwell-based epithelial airway models have been used in diverse applications, ranging from complex co-culture configurations [9] and high-throughput formats [10] in 96-well plates to primary cell models of cystic fibrosis patients and toxicity assessments [11–13].

While transwell inserts become increasingly popular, the inherent static culturing conditions fail to recapitulate dynamic physiological processes, such as fluid flow and mechanical forces encountered *in vivo*. Therefore, advanced microphysiological *in vitro* models were developed to better recapitulate the physiological microenvironment of the human airway system with higher physiological relevance [4]. Since Huh’s microfluidic alveolar lung device in 2010 [14] set the starting point of lung-on-chip platforms, several microphysiological airway systems have been proposed. Developments also include bronchial airway models of varying complexity, ranging from highly relevant primary stem cell-derived systems to epithelial-endothelial coculture models [15, 16].

However, most microphysiological airway cultures are generated in the microfluidic device itself, often requiring laborious protocols with limited throughput. Moreover, many stem-cell-based tissue models were originally developed and optimized in well-plate-based cultures and require several weeks to achieve proper cell differentiation. A direct transfer of these protocols to more complex microphysiological systems is not always feasible.

To overcome this limitation, transwell-based microphysiological systems have been introduced. In such systems, commercially available transwell cultures were inserted in a microfluidic environment to study alveolar epithelium barrier functions [17], the effect of cannabis smoke on bronchial epithelial cells [18], 3D morphogenesis of intestinal epithelial cells [19], and the directional alignment of endothelial cells and myotubes under flow stimulation [20].

Nevertheless, both conventional transwell systems and microphysiological models exhibit substantial imaging limitations. Typically, these systems feature a significant working distance or require imaging through multiple materials. These characteristics impede the capturing of high-resolution images of the tissue layer, making it challenging to resolve details at the single-cell level. Since most tissue models are cultured on the upper surface of the transwell insert, light must pass through the porous membrane, further compromising imaging quality and impeding observations of cellular features at high resolution. Moreover, the routinely used air-liquid interface culture complicates compatibility with high-NA objectives, which typically require liquid interfaces to achieve optimal resolution.

However, imaging tissue samples at high spatiotemporal resolution is crucial for understanding complex cellular processes and for observing cellular dynamic changes in response to perturbations, such as infections. Continuous live-cell imaging could give unprecedented insights into host-pathogen interactions, filling the current knowledge gap and opening new perspectives for the discovery and assessment of novel therapies.

Here, we present a microphysiological system, designed for live-cell, high-resolution imaging of transwell-based airway tissues. This plastics-based device offers seamless transition between air and liquid-phase culturing conditions. The airway epithelium is cultured most of the time under physiological conditions in air, while perfusion of liquid is periodically applied for a short time to enable high-spatiotemporal-resolution imaging through the liquid. The modular platform design allows for transwell insertion and removal at any stage of the experiment, making it compatible with established well plate-based culture protocols and providing the flexibility to select the most suitable insert for on-chip cultivation. Furthermore, the damage-free extraction of inserts facilitates the use of a wide range of downstream analysis methods for in-depth tissue sample analysis. In this study, we mimicked *Pseudomonas aeruginosa* infections of the bronchial epithelium and continuously imaged infection progression in real time at high resolution. We monitored bacterial spreading and invasion of *P. aeruginosa* on the apical epithelial surface and visualized tissue breaching and destruction over time. Overall, these findings demonstrate the capabilities of the system to elucidate key mechanisms of respiratory infections. We anticipate that the modularity of the platform and its compatibility with commercial transwell inserts make it a useful tool for studies of other barrier tissue models.

## 2. Materials and methods

### 2.1. Microfluidic device fabrication

Poly(methyl methacrylate) (PMMA)-based microfluidic devices (25 mm x 75 mm) were fabricated using a computer numerical control (CNC) micro-milling system (Roland SRM-20) as described elsewhere [21]. The chip featured four identical micro-milled channel units of 700 μm height, each with an open well in the center, where a cell-coated transwell was inserted. The bottom of the channel was sealed with a glass coverslip (ref. 10812, ibidi GmbH, Martinsried, Germany) using a biocompatible epoxy glue (MED-302-3M, Epoxy Technology Europe AG, Cham, Switzerland).

A stable and leak-free insertion of the transwell was achieved through a ring-shaped adhesive gasket (GBL620004, Grace Bio-Labs, OR, USA) that tightly sealed the insert. The basolateral side of the transwell was closed with a custom-designed GreenTec Pro-based lid to prevent contamination and minimize media evaporation. Each channel contained inlet (diameter = 0.8 mm) and outlet (diameter = 0.8 mm) openings for liquid perfusion. Additionally, each fluidic unit featured four metal needles for liquid aspiration from the apical compartment of the inserted transwell. The set of aspirating needles consisted of two straight needles and two angled needles (IPS618150, GONANO Dosiertechnik GmbH, Breitstetten, Austria) positioned close to the cell-covered transwell. The fully assembled chip was sterilized before the fluidic experiments by washing with 50% ethanol and UV treatment from both sides for 30 min.

**Figure S1** shows the full experimental setup. Cell-coated transwells were manually inserted into the open well in the center of the chip and sealed by the ring-shaped gasket. The chip was placed inside a microscope stage incubator at 37°C in an atmosphere of 5% CO^2^ and 95% relative humidity. 10 ml glass syringes (ILS, Ilmenau, Germany) were mounted to syringe pumps (neMESYS, Cetoni GmbH, Korbussen, Germany). Syringes were connected via standard Luer-lock syringe-tubing connectors (TE 27 GA 90° bent, APM Technica AG, Heerbrugg, Switzerland), polytetrafluoroethylene (PTFE) tubing (ID 0.3 mm, OD 0.6 mm, Bola GmbH, Grünsfeld, Germany) and metal connecting pieces, obtained from standard Luer-lock syringe-tubing connectors (TE 27 GA 90° bent, APM Technica AG, Heerbrugg, Switzerland), to the inlets of the chip. The inflow rate of the syringe pumps was set to 10 μl min^−1^. We used a peristaltic pump (ref. 78001-18, Ismatec) to aspirate HBSS from the apical channel at a total flow rate of 120 μl min^−1^and re-establish air-liquid-interface (ALI) conditions. Upstream, a peristaltic tubing (Tygon S3 E-LFL, ID 0.27 mm, wall 0.91 mm, Idex Health & Science GmbH, Wertheim, Germany) was connected to each aspirating outlet.

### 2.2. Cell culture

Normal Human Bronchial Epithelial (NHBE) cells were obtained from Lonza and differentiated according to the protocol provided by StemCell Technologies. First, the cells were thawed and grown in a T75 flask using PneumaCult-Ex Plus medium (Stemcell Technologies, Cologne, Germany). Once confluency reached 60–80%, the cells were dissociated using Animal Component-Free (ACF) enzymatic dissociation solution (Stemcell Technologies, Cologne, Germany). Then, the cells were seeded on the bottom side of the permeable membrane (0.4 μm pore size) of 6.5 mm Transwell inserts (Costar, Corning, NY, USA). Cells were seeded at a density of 3.3 × 10^4^ cells/insert and grown at a liquid-liquid interface in a PneumaCult-Ex Plus medium. After the cells reached full confluency, the medium was removed from both sides of the tissue, and the basolateral side was filled with 350 μl PneumaCult-ALI medium to establish an air-liquid interface. The medium was exchanged every second or third day. From an age of 25 days, the apical side was washed once a week with warm HBSS to remove any excessive mucus accumulated on the tissue surface. All experiments were performed with tissues cultured at the ALI for at least 25 days.

### 2.3. Viability assay

Cytotoxicity was assessed by measuring the release of lactate dehydrogenase (LDH) into the culture supernatant using the cytotoxicity detection kit PLUS (Roche, Basel, Switzerland). 100 μl cell-free supernatant, collected from the basolateral compartment, was mixed with 100 μL reaction mixture and added into each well of 96-well flat-bottom polystyrene plates (Costar, Corning, NY, USA). The plates were incubated at 37°C for max. 30 min while being protected from light. The reaction was stopped by adding 50 μl of stop solution, and the absorbance was detected at a wavelength of 492 nm using a multi-detection reader. The LDH positive control was generated by treating the tissue with the lysis buffer from the cytotoxicity kit for 24 h.

### 2.4. Cilia beating frequency measurements

Live cilia beating movies were acquired using the following setup: Nikon CSU-W1 microscope, equipped with a Plan Apo Lambda 4x/0.13 objective, at 37°C in an atmosphere of 5% CO_2_ and 95% relative humidity. The cilia beating frequency was expressed in frequency units (Hz). For each movie, 250 images were captured at a high frame rate (200 frames per second). Cilia beating frequency was then measured using Cilia-X software (Epithelix, Geneva, Switzerland).

### 2.5. TEER measurements

Transepithelial electrical resistance (TEER) of the tissues was measured by an EVOM3 Volt-Ohm resistance meter (World Precision Instruments, Berlin, Germany), equipped with STX2 chopstick electrodes (World Precision Instruments). After mucus washing, the tissue was submerged in 37°C warm HBSS (350 μl apical and 1 ml basolateral) for TEER measurements. Reported TEER values were derived by normalizing each endpoint measurement with respect to its initial measurement value.

### 2.6. Immunocytochemistry

Tissues were fixed by adding 10% formalin to both apical and basolateral compartments of the transwell for 45 min at RT. Once fixed, the formalin was removed, and tissues were washed with HBSS containing 0.05% NaN_3_ and stored at 4°C until staining. The cell-covered membrane was manually cut out, and tissues were permeabilized with 0.5% Triton-X (Sigma-Aldrich, Buchs, Switzerland) diluted in HBSS for 20 min at RT. After one round of washing in HBSS with 0.05% NaN_3_, tissues were stained: diluted (1:10,000) CellMask Deep Red (C10046, ThermoFisher Scientific, Waltham, MA, USA) was used as a marker of the plasma membrane, a 1:200-diluted anti-ZO-1 antibody (MA3-39100-A555, ThermoFisher Scientific, Waltham, MA, USA) was used for staining tight junctions, and cell nuclei were stained with 1 μM DAPI 33342 (D9542, Sigma-Aldrich, Buchs, Switzerland). The staining was conducted for 4 hours at RT. After washing in HBSS with 0.05% NaN_3_, the membrane was mounted onto a microscope slide (25 mm × 75 mm), covered with one drop of Mowiol, and sealed with a cover slip (No. 1.5H, 22 mm × 22 mm, Marienfeld GmbH, Lauda-Königshofen, Germany). The embedded samples were cured for 48 hours at RT before imaging. The images were acquired with a Nikon CSU-1 spinning-disc confocal microscope (Nikon, Egg, Switzerland) using a 100× objective (NA 1.4, WD 0.13 mm).

### 2.7. Liquid-liquid interface assay

Fully mature airway tissues were used for liquid-liquid interface (LLI) tests after at least 25 days of ALI differentiation. First, medium from the basolateral compartment was collected to establish a baseline measurement for the tissue viability assay (see 2.3). TEER was measured to determine the initial barrier function. To establish an LLI, the apical surface of the insert was filled with 600 μl of synthetic cystic fibrosis sputum medium (SCFM), Phosphate saline buffer (PBS), and Hank’s balanced salt solution (HBSS). After 24 h, basolateral medium was collected for the viability assay, and the barrier function was evaluated using the TEER assay. Both TEER values and viability data were reported as relative values of endpoint (t24h) to initial (t0h) conditions.

### 2.8. Tissue infection

*Pseudomonas aeruginosa* were cultured in a synthetic cystic-fibrosis medium (SCFM) at 37°C continuous shaking. Once the culture reached a stationary phase, it was diluted at a 1:100 ratio in SCFM and grown for 3 hours under the same conditions. After growth, bacteria were washed, and their concentration was adjusted to an optical density (OD600) of 1.0 using phosphate-buffered saline (PBS). Lung tissues were infected with 2 μl of the bacterial suspension using an inoculum size of 0.5 × 10^6^ CFU/ml.

### 2.9. Live imaging of on-chip infection with P. aeruginosa

The tissues were stained with 2 μg ml^-1^ CellMask™ Deep Red plasma membrane stain (ThermoFisher Scientific, Waltham, MA, USA) for at least 24 h before imaging. The imaging setup included a Nikon CSU-W1 equipped with a 2x Hamamatsu ORCA-Fusion camera (sCMOS) and CFP Plan Apo Lambda 10x/0.45 and CFI Apo LWD Lambda S 40x/1.15 water objective. Images were analyzed using NIS-Elements General Analysis software and Imaris Imaging Analysis software.

### 2.10. Statistical analysis

Statistical analysis was performed using GraphPad Prism. Differences between experimental groups were assessed using a two-tailed unpaired t-test with Welch’s correction (*, p < 0.05; **, p < 0.005; ***, p < 0.0005). All results are shown as mean values ± SEM.

## 3. Results

### 3.1. Concept of the microphysiological transwell-based platform

We developed a microphysiological platform to address the imaging limitations of conventional transwell cultures and to expand the experimental capabilities, as illustrated through a side-by-side comparison in **Figure 1**. We inverted the orientation of the tissue layer of conventional transwell cultures by growing the airway epithelium at the bottom side of the permeable membrane, which enables direct access of the microscope objective and prevents image obstruction by the porous membrane. The inverted setup in the microfluidic device allows for non-invasive collection of medium from the basolateral side of the transwells for downstream assays without perturbing live imaging, whereas conventional transwell systems require a removal of the inserts from the well plate for basolateral medium sampling. We also significantly reduced the distance between the cell layer and the microscope objective from approximately 1 mm with conventional transwell inserts to 0.30 mm in the microphysiological platform. This shorter working distance provides flexibility for imaging of the tissue with a wide range of objective lenses with magnifications of up to 60x.

**Figure 1.**
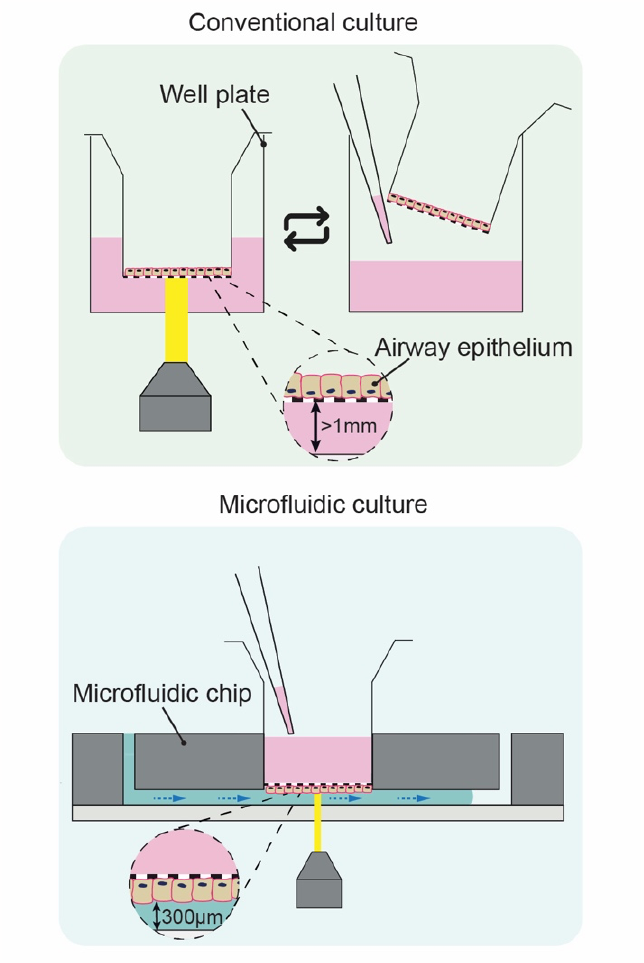
Side-by-side comparison of conventional transwell cultures and the microphysiological transwell-based platform. In the microfluidic setup, airway tissues are cultured on the bottom side of the membrane, while conventional transwell cultures typically feature the tissue on top of the membrane. For the imaging assay, transwells are inserted into the microfluidic chip at a working distance of 300 μm from the microscope objective. The chip is fluidically connected to establish a liquid-liquid interface for high-resolution imaging. In contrast, conventional transwells exhibit a working distance of 1 mm and require imaging through the semipermeable membrane, which impedes high-resolution imaging. In the microfluidic setup, basolateral medium can be sampled continuously from the chip during imaging, whereas the conventional setup requires to interrupt the imaging process for sampling.

Furthermore, the microfluidic design allows for infusion of controlled volumes of liquids, enabling immersion of the tissue surface in medium, which is essential for the application of high-resolution imaging.

### 3.2. Device design and operation

We employed human stem cell-derived bronchial epithelial tissues grown on commercial transwell inserts of 0.4 μm pore size as our experimental airway model. This model exhibits a pseudostratified epithelium with in vivo-like morphology and mucociliary clearance upon differentiation, as reported previously [3, 4].

First, we evaluated the compatibility of the system with high-resolution imaging by testing the effect of a non-physiological liquid-liquid interface (LLI) on the airway model. We exposed the apical surface of the airway epithelium to synthetic cystic fibrosis sputum medium (SCFM) [22, 23], phosphate saline buffer (PBS), and Hank’s balanced salt solution (HBSS). SCFM is typically used to mimic sputum of cystic fibrosis patients, whereas PBS and HBSS are commonly used in airway infection assays. Notably, we aimed at establishing conditions as similar to in-vivo physiology as possible by avoiding nutrient-rich maintenance media, such as PneumaCult-ALI medium.

We found no significant viability variations of airway tissues across all tested LLI conditions after 24 hours. This finding was supported by the low LDH release, which was similar to that of the negative control cultured at an air-liquid interface (ALI) (**Figure S2a**). Nevertheless, the barrier integrity of the tissue models under SCFM, PBS, and HBSS conditions was substantially compromised, as evidenced by a significant reduction in transepithelial resistance (TEER) over the culture period (**Figure S2b**).

Prompted by the disruptive effect of prolonged LLI culturing on the airway model, we sought to develop a device to maintain airway models under physiologically relevant ALI conditions while enabling short-term high-resolution imaging sessions under LLI conditions. We fabricated the device using inert, biocompatible polymethyl methacrylate (PMMA) in a standard microscopy slide format. The use of standard thermoplastic material alleviates the issue of non-specific compound absorption, which is characteristic for PDMS-based organ-on-a-chip platforms [24]. Each device featured four identical culture wells for hosting commercially available transwell inserts. The bottom face of the device, featuring the fluidic structures, was sealed with a glass coverslip. We utilized a 3D-printed lid to close and protect the basolateral compartments of the transwells during the on-chip culturing (**Figure 2a-b**). We ensured a seamless transwell insertion by integrating a ring-shaped sealing gasket around the culture well (**Figure 2c**). Once a transwell was inserted, the gasket provided a robust seal for long-term, leakage-free perfusion of the channel.

**Figure 2.**
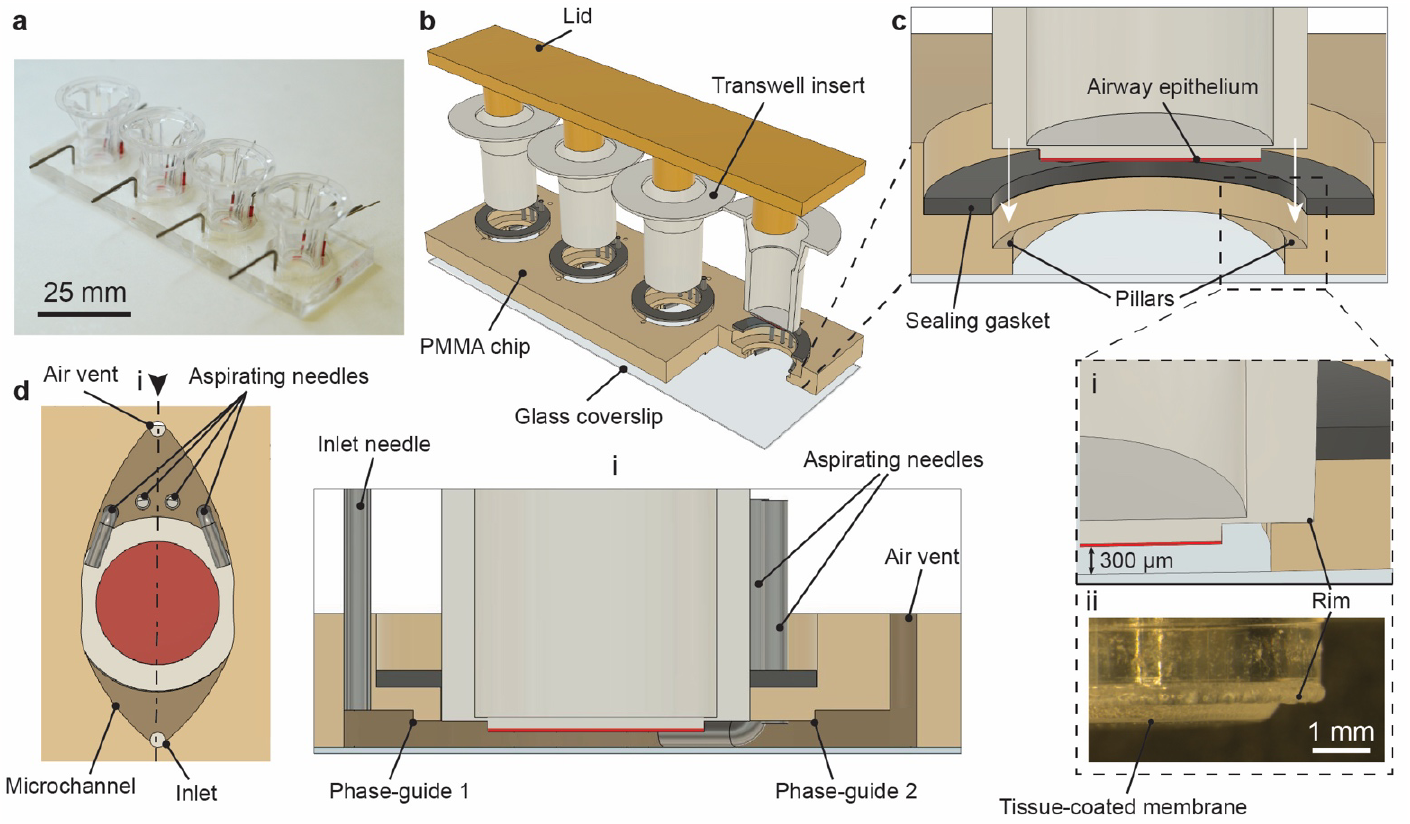
Microphysiological platform for culturing and imaging of transwell-based airway models. **(a)** Photograph of the fully assembled device. **(b)** A schematic representation of the PMMA-based device that contains four culture units, each accommodating one transwell insert. A glass coverslip seals the chip from the bottom, while the transwells are closed with a 3D-printed lid from the top. **(c)** Insertion of a transwell into the culture unit of the chip. The plastic rim lands on top of the protruding pillar structures. A fully inserted transwell (i) is fluidically sealed from the sides by a ring-shaped silicone gasket, and the tissue-coated membrane (ii) is positioned 300 μm above the glass coverslip. **(d)** Top-view schematics of the microchannel with inlet, air vent, and tips of aspirating needles. The tissue coated membrane is shown in red. Cross-sectional view (i) shows the phase-guide structures and aspirating needles in close proximity to the tissue-coated membrane (red) in the microchannel.

Pillar-like protruding edges were introduced at the sides of each culture well to define the position of the transwell at a predetermined distance from the coverslip surface, ensuring a consistent working distance during imaging. To prevent tissue damage, we designed the pillars to only overlap with the plastic rim around the permeable membrane on the insert. As a result, the tissue barrier was positioned at a vertical distance of 300 μm from the glass coverslip, as shown in the close-up in **Figure 2c**.

We established a liquid interface at the apical surface of the tissue by filling the channel with HBSS through an inlet needle at a flow rate of 10 μl min^−1^ for 5 min (**Figure 2d, Movie S1**).

Bubble-free perfusion under the insert was achieved by introducing two 100-μm-high phase-guides that facilitated uniform channel filling. Once HBSS entirely covered the apical surface of the tissue, we used a 40x water immersion objective (NA = 1.15) to image the cells at high magnification. After the imaging has been completed, we re-established the air interface at the apical surface of the tissue: we sealed the air vent and removed HBSS from the channel through peristaltic pump-driven aspiration through four aspirating needles (**Figure 2d**). Aspirating needles were positioned in immediate proximity to the airway epithelium to ensure full removal of HBSS below its apical face. Each aspirating needle, set to withdraw at 30 μl min^−1^, achieved a combined flow rate of 120 μl min^−1^, resulting in complete HBSS removal within 30 seconds (as demonstrated in **Movie S1**).

### 3.3. Modular culture of transwell-based airway models

The implementation of the microfluidic transwell-based assay is shown in **Figure 3**. We devised the assay in a modular format, providing flexibility to insert and remove the transwells at any desired time. In the first step, we seeded NHBE stem cells on the bottom side of the transwell inserts (1)and cultured them at a liquid-liquid interface until full confluency was reached (2). Subsequently, the tissue was differentiated at an air-liquid interface for at least 25 days (3). The off-chip culturing offers the possibility of integrating complex tissue models into the platform, which often require growing over several weeks under well-defined conditions. Moreover, standardized protocols allow for generation of large tissue batches and for picking only certain transwells for experimentation after conducting simple, non-invasive assays, such as TEER measurements (4).

**Figure 3.**
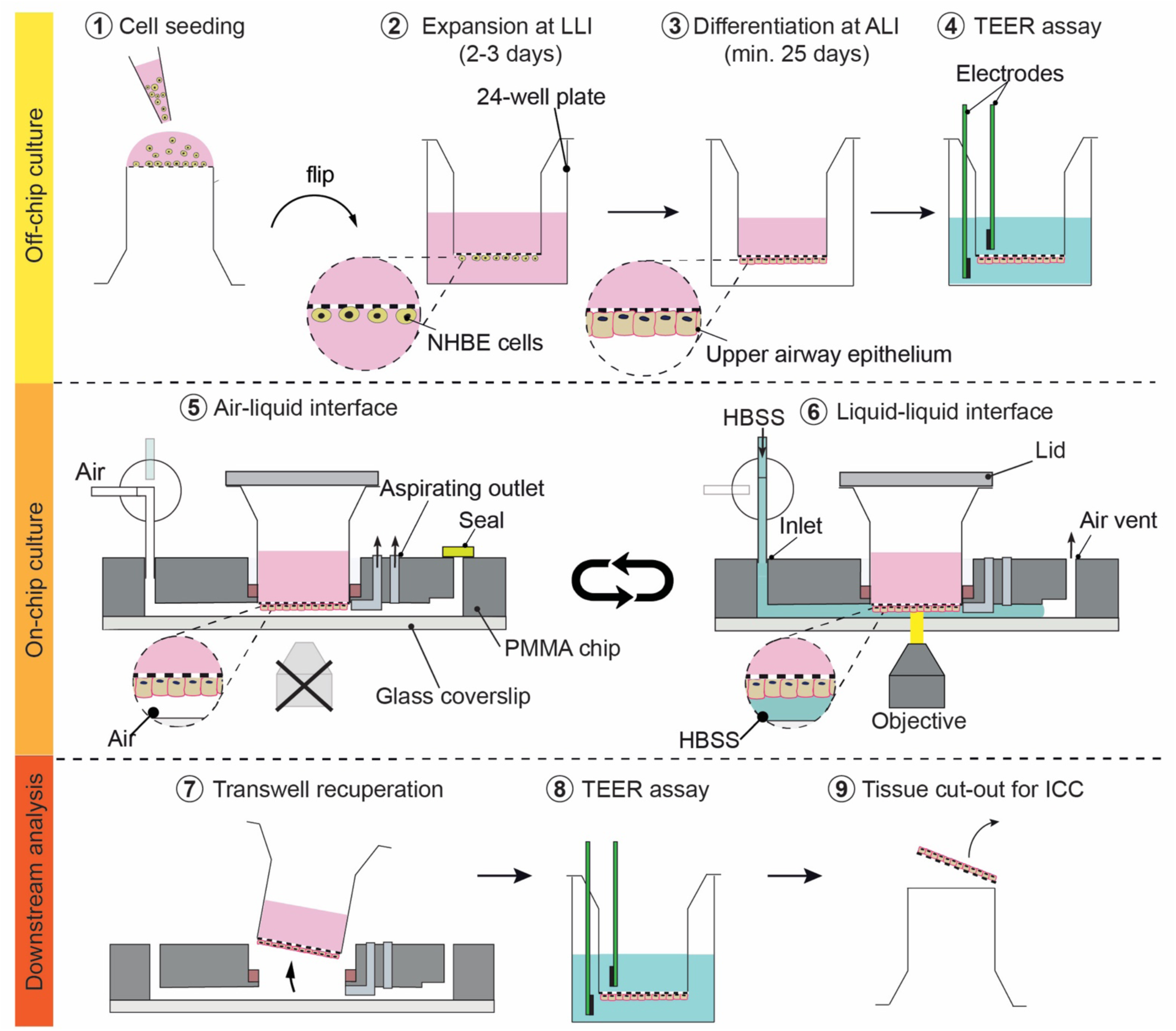
Experimental workflow of the modular transwell-based airway epithelium culture. Off-chip culture: 1,2) NHBE cells are seeded on the bottom surface of the PET insert and grown at the LLI. 3) Once the cells are fully confluent, an ALI is established for a minimum of 25 days to facilitate differentiation of the cell layer into pseudostratified epithelium. 4) Transwells with fully differentiated tissue models are assessed by TEER and are then selected for insertion in the microfluidic device. On-chip culture: 5) Airway tissues in the microphysiological platform are cultured at a physiologically relevant ALI. 6) An LLI is established by infusing HBSS into the microchannel. The immersed state is utilized for high-resolution imaging with a high-NA objective. Once imaging has been completed, an ALI is re-established by withdrawing HBSS through the aspirating outlets. 7) The transwell is recuperated from the chip after culturing by alternating between ALI and LLI, then subjected to downstream analysis by 8) a TEER assay and 9) fixation and immunocytochemical staining.

Next, we inserted the selected transwells into the culture well of the microphysiological platform and maintained them under ALI conditions until starting the imaging process (5). For high-resolution imaging, we connected an inlet needle to a HBSS-filled syringe and introduced the buffer solution to the apical side of the airway tissue, creating optimal imaging conditions at the LLI (6). Upon completion of the imaging protocol, we re-established ALI conditions for tissue maintenance until the next imaging time point. During liquid aspiration from the channel, the perfusate was collected non-invasively from the system for secretome analysis of the apical tissue face. After the experiment, we recuperated the intact tissue model from the platform (7) to perform in-depth downstream analysis at tissue and single-cell level (8, 9).

### 3.4. Preservation of barrier integrity and viability of airway models during alternation of air-liquid and liquid-liquid interface conditions

As described previously, the transition between ALI and LLI conditions is critical to reproducing the human airways’ physiological *in vivo* environment while enabling high-resolution imaging. Therefore, we examined the effect of recurrent transitions between ALI and LLI conditions on the airway models during a 6.5-hour-long on-chip culture (**Figure 4a**). First, we filled the channel with HBSS and submerged the apical surface of the tissue in liquid for 30 minutes. We then removed HBSS via pump-driven aspiration, thus re-establishing an ALI for 90 minutes. We repeated this process 3-times before recuperating the transwell from the chip for analyzing the airway model. As a control, we utilized airway tissues maintained continuously at an ALI in the microphysiological chip for 6.5 hours.

**Figure 4.**
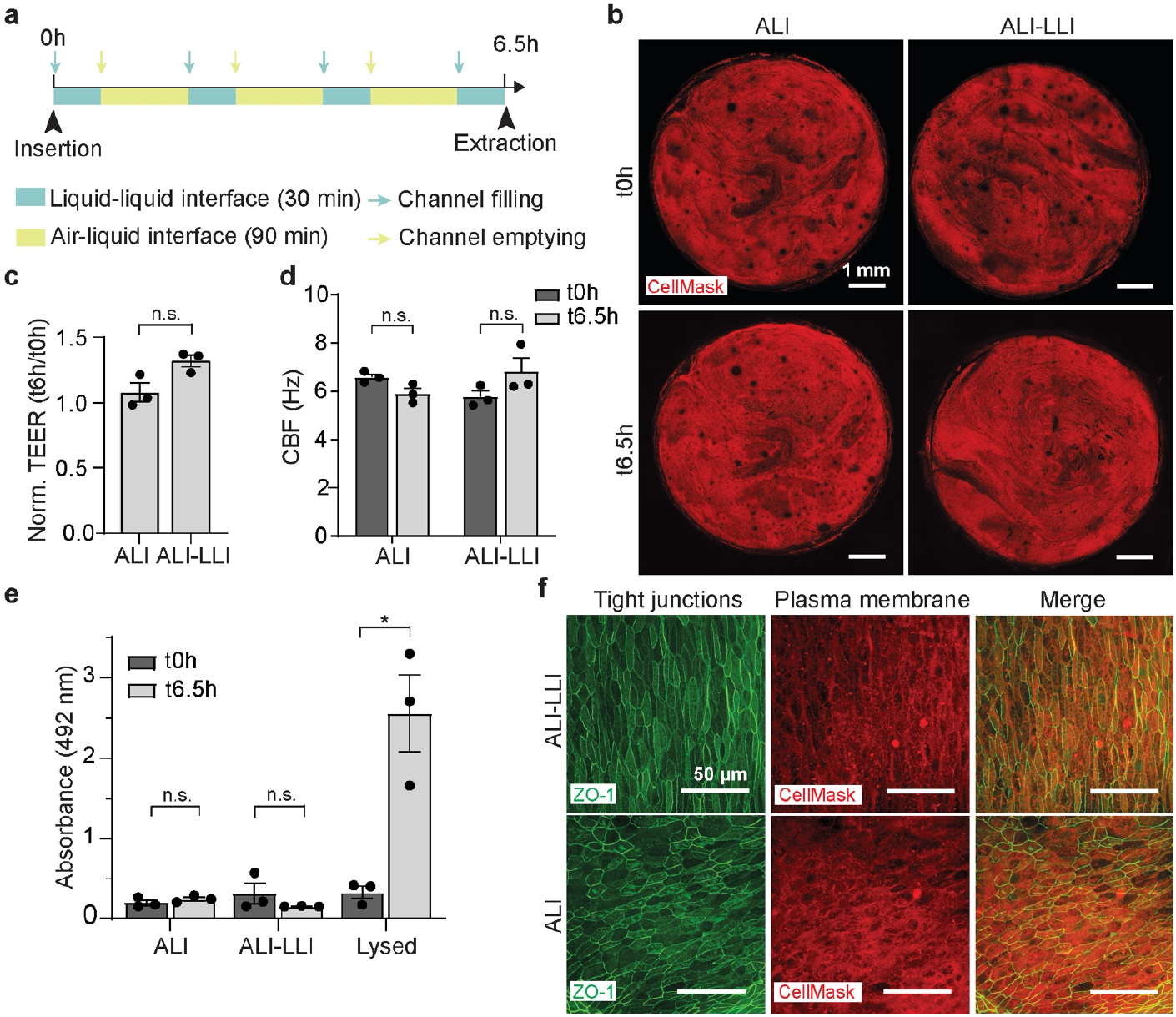
Preservation of barrier integrity and viability of airway model after ALI-LLI alternation on-chip. **(a)** The experimental timeline included transwell insertion in the chip (t0h), periodic alternation between air-liquid interface and liquid-liquid interface (ALI-LLI) on-chip for 6.5 h, and transwell recuperation for downstream analysis (t6.5h). **(b)** Overview of the entire airway tissue cultured for 6.5 h under either ALI or ALI-LLI conditions on chip. **(c)** Normalized epithelial barrier integrity (TEER), **(d)** Measurements of cilia beating frequencies (CBF), **(e)** Tissue viability (LDH release) of transwells cultured under ALI and ALI-LLI conditions on chip. The positive control group was cultured with an addition of lysis buffer. **(f)** Tight junctions (ZO-1) and cell membranes (CellMask) of airway epithelium cultured under ALI and ALI-LLI conditions on chip. Mean values ± SEM, n = 3 transwell inserts per condition.

We confirmed that the tissue integrity was not affected by multiple rounds of ALI-LLI alternations, as shown by images of the entire tissue surface obtained at the beginning (t0h) and the end of the experiment (t6.5h, **Figure 4b**). Moreover, alternation did not affect the barrier integrity of the airway model, as TEER measurements did not show any differences between ALI alone and alternating ALI-LLI conditions (**Figure 4c**). These results also confirmed that the mechanical process of inserting and recuperating the insert from the chip did not affect tissue integrity and tightness.

We further tested the effect of ALI-LLI alternation on the mucociliary clearance activity of the airway tissue. **Figure 4d** shows no significant difference in cilia beating frequency (CBF) upon ALI-LLI alternation, indicating active and coordinated beating of cilia on the tissue surface. The cytotoxicity measurements corroborated tissue viability, as there was no significant increase in LDH release upon ALI-LLI alternation (**Figure 4e**). Immunocytochemical (ICC) staining of tight junctions (ZO-1) and the plasma membrane (CellMask) showed that the structure of the epithelium was well preserved (**Figure 4f)**, proving the suitability of the device for imaging assays at the epithelial barrier.

### 3.1. High-resolution imaging of airway model infection under interface alternation in the chip

We sought to demonstrate the device’s imaging capabilities by emulating airway epithelium infection by *Pseudomonas aeruginosa* under alternating ALI-LLI conditions. It has been reported that the critical steps in the *P. aeruginosa* infection process, such as widespread bacterial internalization and tissue breaching, occur at approximately 15 hours post-infection in the model employed here [23]. Therefore, we designed our experimental timeline to commence on-chip culture and high-resolution imaging at 13 hours post infection, continuing until 19 hours post infection (**Figure 5a**). This approach enabled us to capture the most pivotal stages of *P. aeruginosa* infections at high spatiotemporal resolution.

**Figure 5.**
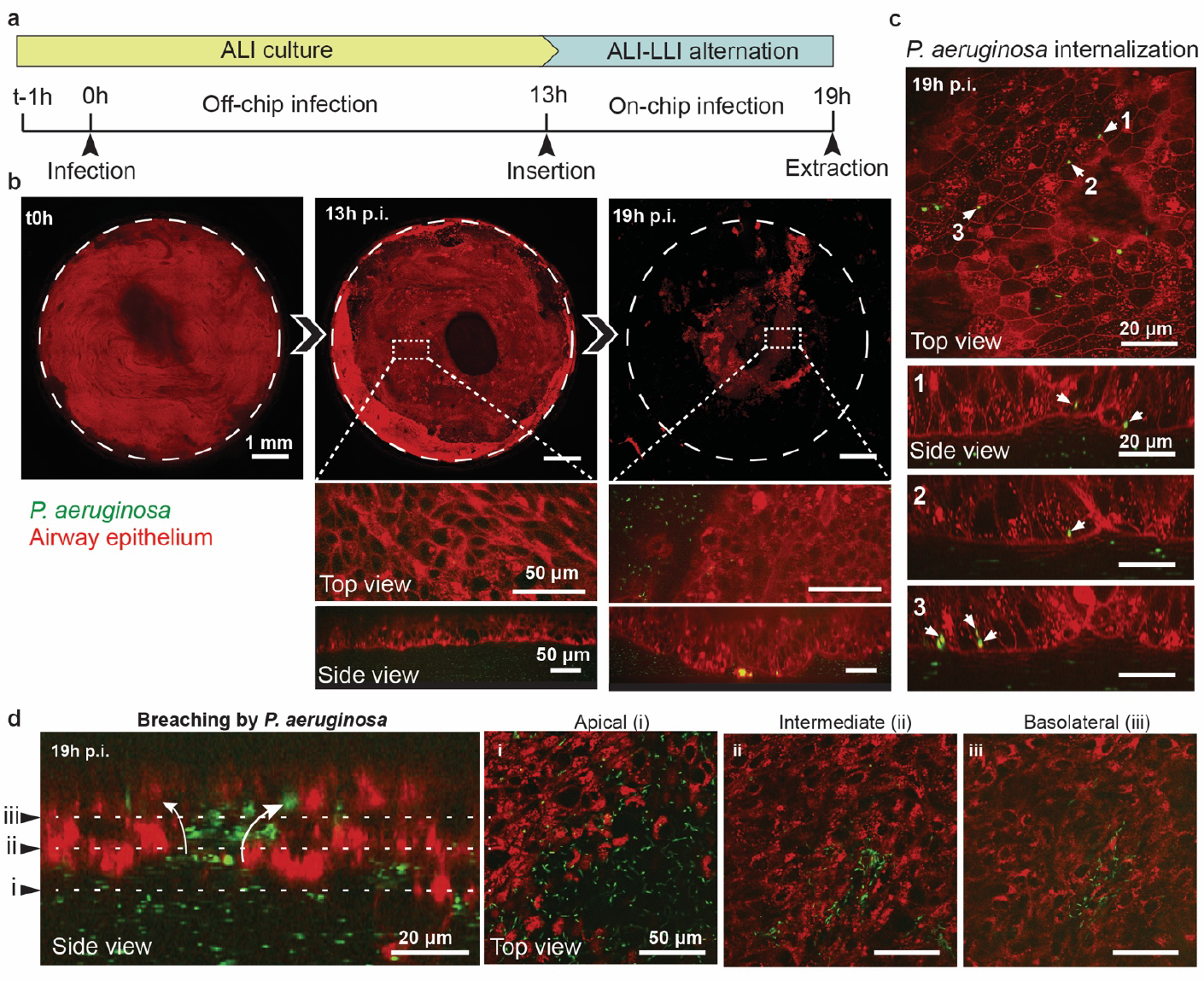
Live microscopy images of an upper airway epithelium infection by *Pseudomonas aeruginosa*. **(a)** Experimental workflow. Airway organoids are inoculated with bacteria and incubated for 13 h in a well plate at an ALI. The transwell is inserted in the chip at 13 h post-infection (p.i.) and subjected to alternations of ALI-LLI for 6 h until recuperation for downstream analysis. **(b)** Initially intact airway tissues (t0h) show localized damages at 13 h p.i. and lose tissue integrity by 19 h p.i. High-resolution images also unveil abrupt changes in the tissue barrier morphology (top view) and flatness (side view) during infection progression. **(c)** High-resolution images capturing widespread *P. aeruginosa* internalization within airway epithelial cells. **(d)** Tissue breaching by *P. aeruginosa*, with large bacterial colonies at the apical surface (i) that progress further into intermediate (ii) and basolateral (iii) layers of the epithelium.

First, we focused on overall tissue damage at the onset of the infection. At 13 hours post infection, we observed localized destruction of the epithelial barrier at the insert’s edges, in contrast to the intact tissue observed before infection (**Figure 5b, Movie S2**). Interestingly, the central region of the insert featured a largely intact cellular barrier, as evidenced by its healthy tissue morphology and uniform thickness (**Figure 5b**). However, at 19 hours post infection, the tissue barrier was significantly compromised, leaving only small areas of tissue still attached to the insert. Accordingly, high-resolution images at this time-point revealed major deformations in the airway epithelium morphology.

Live images of the infected airway tissues revealed the internalization of *P. aeruginosa* within the airway epithelium (**Figure 5c, Movie S3**). We simultaneously monitored multiple regions of the tissue surface and observed several epithelial cells containing single internalized bacteria (1, 2), while other regions contained a few bacteria in a single cell (3), potentially indicating replication within the cell body. Live images also captured the formation of breaching points within the tissue barrier, which constitutes a hallmark of *P. aeruginosa* infection, as disruption of the barrier leads to a rapid basolateral invasion of the bacteria and full tissue destruction [23, 25]. As shown in **Figure 5d**, we captured the phase of the infection breaching through the entire thickness of the tissue. We observed that *P. aeruginosa* bacteria densely colonized the apical surface (i) and breached the ciliated barrier for invasion of the underlying intermediate (ii) and basolateral (iii) layers. Notably, in contrast to these real-time high-magnification (40x) observations of *P. aeruginosa* internalization, breaching events in airway models have previously only been visualized using fixed tissue samples.

## 4. Discussion and conclusion

Studying tissue samples at a microscopic scale is essential for uncovering the underlying mechanisms of cellular processes under both healthy and diseased conditions. However, tools for investigating transwell-based organoid cultures at high spatiotemporal resolution are scarce. Therefore, we developed a microphysiological system tailored for transwell-based airway tissues that enables real-time high-resolution imaging. To demonstrate the system’s capabilities, we emulated a *Pseudomonas aeruginosa* infection of the bronchial epithelium, which showcases the potential to obtain novel insights into airway infection dynamics.

Airway epithelial tissue is naturally exposed to an air interface, serving as the first line of defense against airborne pathogens [26, 27]. To recapitulate this *in vivo* environment in an *in vitro* setting, transwell inserts are commonly employed to grow tissue barriers on top of a semipermeable membrane, which is exposed to ambient air on the apical tissue side [10, 28]. Protocols have been developed to culture the air-exposed epithelial cells at the bottom side of the membrane to improve imaging conditions and to eliminate image obstruction by the semipermeable membrane [29,30]. Despite these improvements, the working distance in conventional transwell systems is still not compatible with high-resolution imaging of the cellular barrier. Additionally, applying physiological air-liquid interface conditions while using high-resolution optical microscopy poses massive challenges, as the cellular microenvironment of the airway tissue must be adapted to immersion conditions to meet the requirements of high-NA objectives.

To address the limitations described above, we evaluated the compatibility of ALI-based transwell cultures with high-resolution imaging by testing how liquid interfaces affect barrier tightness and viability of bronchial epithelial tissues. We observed decreased barrier tightness under all liquid-phase conditions (**Figure S2**), suggesting potential cellular changes or damage to the epithelial barrier over time. Based on these results, we concluded that a physiologically relevant air–liquid interface is critical for maintaining a healthy airway barrier in our transwell-based models. To still enable high-resolution imaging through a liquid interface, we devised a microphysiological system that alternately switches between air-liquid and liquid-liquid interface conditions (**Figure 3**). To demonstrate the high-resolution imaging capabilities of the developed system, we show a real-time visualization of a *P. aeruginosa* infection in bronchial epithelial tissue (**Figure 5, Movie S2-S3**), a critical clinical pathogen with remarkable intrinsic resistance to commonly used antibiotics [31–33]. A recent study on upper airway infections of *P. aeruginosa* has brought new insights into the underlying infection mechanism of the pathogen [23]. Here, we used a transwell-based organoid model from the same cellular source, however, we cultured the organoids in an inverted orientation, i.e., at the bottom side of the porous membrane. The developed device holds the potential to further advance our understanding of key infection events, including intracellular bacteria activity and tissue breaching as well as host-pathogen interactions governing *P. aeruginosa* infections.

Owing to the high spatiotemporal-resolution imaging needed to study *P. aeruginosa* infections within the scope of this work, the platform exhibits a comparatively low throughput, which represents a limitation of the system. We envision, however, that the platform can be adapted to higher throughput, for example, by 1) integrating non-imaging-based techniques for real-time tissue monitoring (e.g., on-chip-TEER sensor), 2) modifying the current design of the PMMA chip to accommodate 96-well format transwell inserts.

The developed platform could prove valuable in addressing a wide range of biological questions. The use of standard plastic materials and the realization of fluid flow, which can be implemented on both sides of the tissue barrier, will enable to recapitulate in vivo-like dosing profiles for administration of antibiotics in pharmacokinetic studies [34–36]. Combined with imaging and secretome analysis, the platform will enable to draw conclusions on pharmacodynamic effects, ultimately helping to improve treatment concepts and strategies. We foresee that the developed microphysiological system will open new avenues by providing high-resolution optical access to a variety of transwell-based barrier tissue models, such as bladder, placenta, or blood-brain barrier, and in particular to those that are routinely cultured at an air liquid interface.

## Acknowledgements

Authors are grateful for the support of the Imaging Core Facility (IMCF) at Biozentrum, University of Basel, as well as of the Single Cell Facility (SCF) of D-BSSE, ETH Zürich. This work was financially supported by the Swiss National Science Foundation (SNSF) within the framework of the National Competence Center in Research “AntiResist”, New approaches to combat antibiotic-resistant bacteria, under contract number 51NF40_180541.

