## Supplementary Figures for "Transwell-based microphysiological platform for high-resolution imaging of airway tissues"

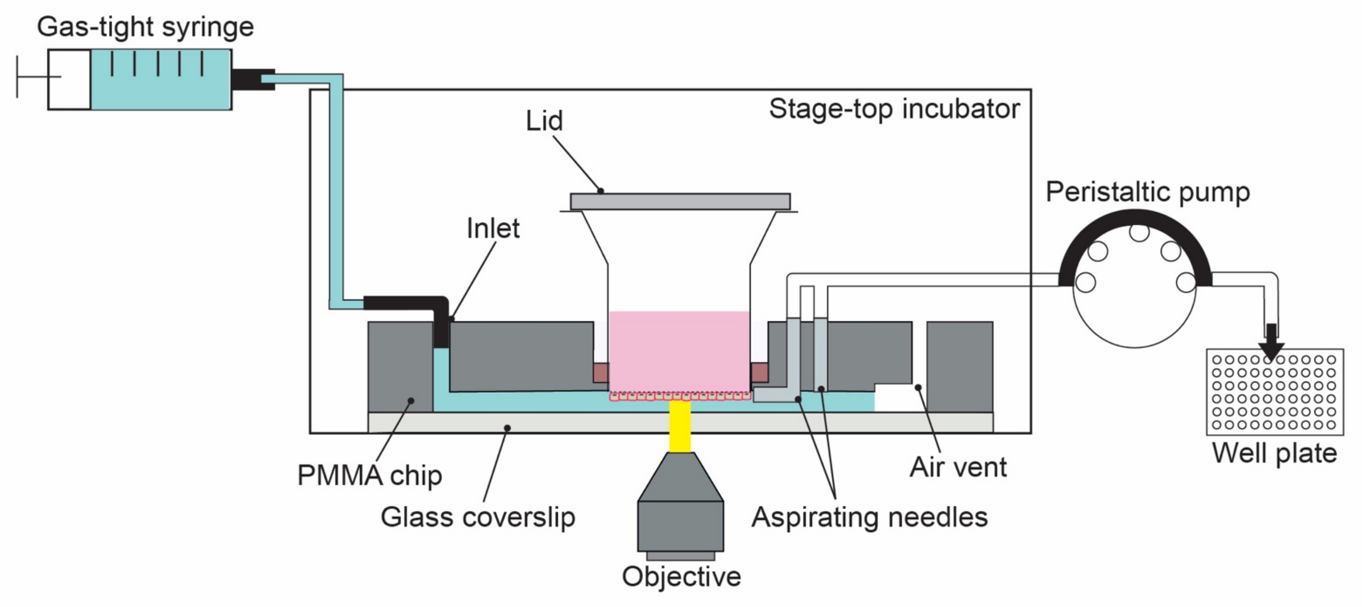


**Supplementary Figure 1. Schematic of the full experimental setup for culturing and imaging of transwell-insert based airway models in the microphysiological platform**. A syringe pump was used to fill the microchannel with HBSS and to establish LLI conditions, and a peristaltic pump is utilized to collect HBSS from the microchannel and to establish ALI conditions. The device was placed on top of an inverted microscope stage, enclosed in a standard commercial stage-top incubator at 37°C, and 5% $CO_2$.

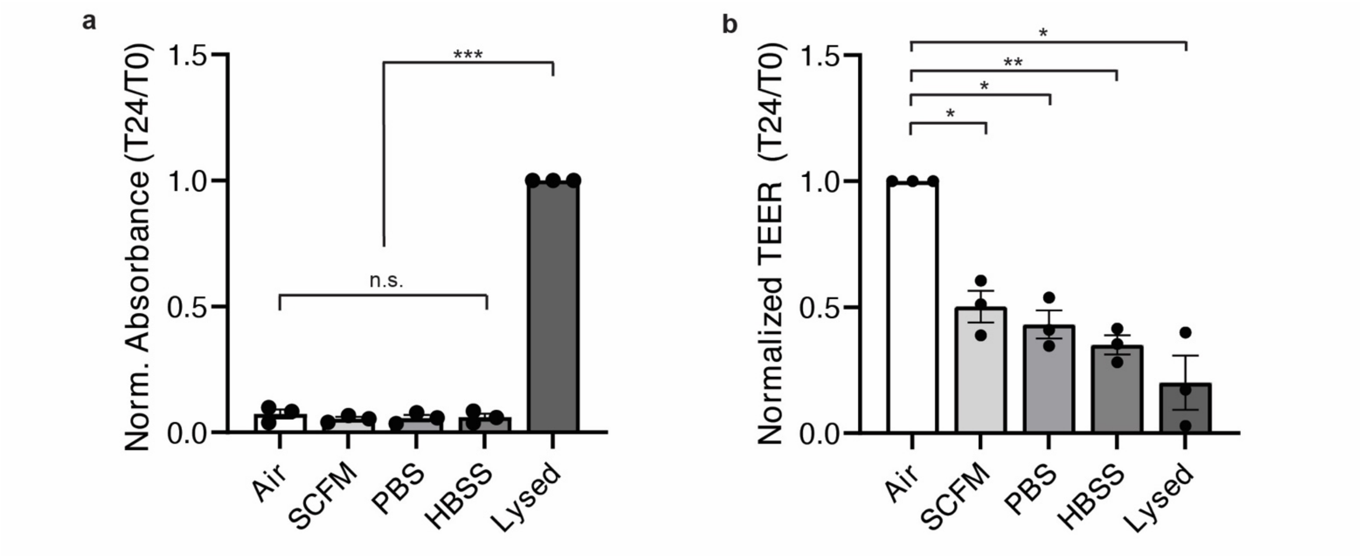


**Supplementary Figure 2. 24h-long liquid-liquid interface assay of upper airway cultures on transwell inserts. (a)** Normalized tissue viability based on LDH release (measured as absorbance). **(b)** Normalized epithelial barrier integrity (TEER). Mean values ± SEM, n = 3 transwell inserts per condition.

**Supplementary Movie 1. Demonstration of the transition between LLI and ALI in the microphysiological platform. HBSS was stained with red amaranth dye for improved visualization of the flow.** (1) Channel filling at 10 µl $min^{-1}$ for 5 min (the movie is sped up 2×). (2) Channel emptying at 120 µl $min^{-1}$ (actual speed)

**Supplementary Movie 2. Live imaging of on-chip infection.** The apical surface of the tissue (membrane: CellMask DeepRed) covered with both motile and adherent *P. aeruginosa* bacteria (mNeonGreen) that started to infect epithelial cells.

**Supplementary Movie 3. 3D animation of the infected airway tissue reconstructed from multiple Z-stacks of live microscopy images.** At 13h p.i. airway tissue (membrane: CellMask DeepRed, nuclei: Hoechst) shows internalized *P. aeruginosa* bacteria in-between apical and basolateral compartments of the tissue.
